# Reconciling Inherent Interfacial Compatibility Conflict Enhances Adhesive Infiltration and Resolves Dentin Bonding Durability

**DOI:** 10.1101/2021.02.09.430396

**Authors:** Qiaojie Luo, Yadong Chen, Jiajia Xu, Chang Shu, Zimeng Li, Weipu Zhu, Youqing Shen, Xiaodong Li

## Abstract

Wet bonding is a basic technique used daily in clinics for tooth-restoration fixation. However, only 50% of the bonding lasts more than 5 years and thus patients must visit the dentists repeatedly. This is attributed to the limited infiltration of adhesives into the demineralized dentin (DD) matrix during wet-bonding. Herein, we show that reconciling interfacial compatibility conflict between the DD matrix and the critical hydrophobic adhesive molecules via hydrophobizing the DD matrix enables the adhesives to thoroughly infiltrate and uniformly distribute within the DD matrix. Thus, the bonding of the hydrophobic DD matrix using commercial dentin adhesives achieves the bonding strength 2-6 times higher than that of the non-treated DD matrix. When a hydrophobic adhesive is applied on the hydrophobic DD matrix, a flawless hybrid layer is produced as observed by nanoleakage investigation. A long-term bonding strength comes up to 7.3 fold that of the control group and very importantly, with no attenuation after 12 months. This study clarifies the basic cause of poor wet-bonding durability and demonstrates a paradigm in adhesive dentistry to overcome the long-existing bonding durability problem associated with inadequate adhesive infiltration into the DD matrix. This provides a new angle of view to resolve clinical dentin bonding durability problem and will significantly promote adhesive dentistry.

**Highlights:**

1. Inherent interfacial compatibility conflict between demineralized dentin matrix and hydrophobic molecules of dentin adhesives is the basic cause for the dentin bonding durability problem.
2. Reconciling the interfacial compatibility conflict markedly facilitates adhesive infiltration in the demineralized dentin matrix and greatly enhances bonding effectiveness.
3. High interfacial compatibility produces a flawless hybrid layer and ideal bonding effectiveness and durability.

**Graphical Abstract:** For wet bonding, poor infiltration of adhesives within the DD matrix inevitably produces numerous defects throughout the hybrid layer, which always leads to the failure of restoration. Via hydrophobizing the DD matrix, reconciling interfacial compatibility conflict between the DD matrix and the hydrophobic adhesive monomers overcomes durability problems associated with the infiltration of adhesives into the DD matrix producing a flawless hybrid layer and providing ideal bonding effectiveness and durability.

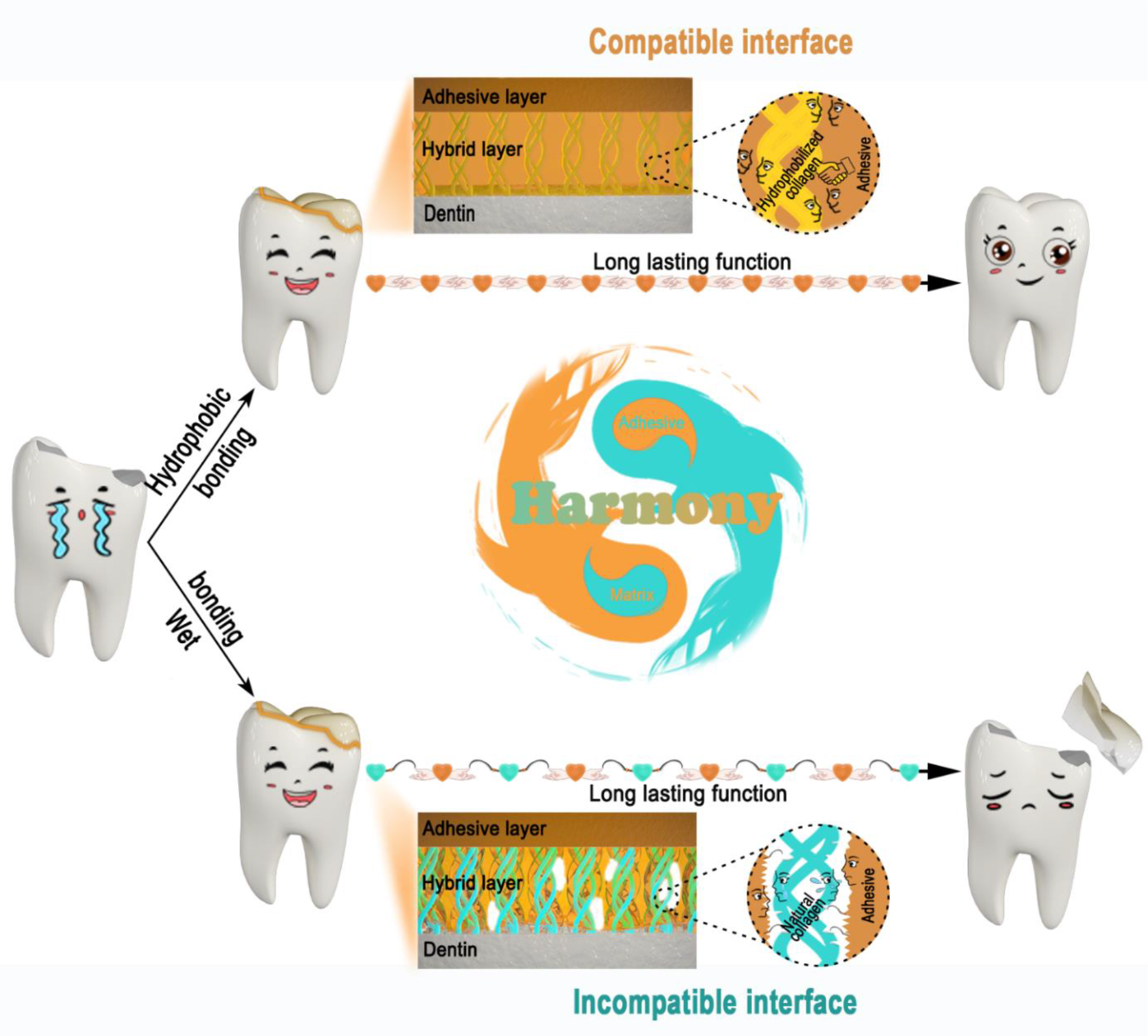

## 1. Introduction

Bonding is basic technique used daily in clinics for tooth-restoration fixation. Contemporary adhesive dentistry is based on wet bonding technique. That is, the DD surface is kept water full to prevent collapse of the DD matrix and provides space for infiltration of the adhesives [1]. The adhesives generally consist of such hydrophobic components as bisphenol A glycidyl methacrylate (Bis-GMA), triethylene glycol dimethacrylate (TEGDMA), and the initiator, camphoroquinone (CQ) and amphiphilic monomers such as hydroxyethyl methacrylate (HEMA) and 2-hydroxypropyl methacrylate (HPMA) as surfactants for the hydrophobic molecules [2, 3]. During wet bonding, the adhesive monomers penetrate into the DD matrix and subsequently polymerize forming the dentin-restoration interface, termed as hybrid layer [4]. Unfortunately, the clinical lifespan of restorations is limited; about half of the restored teeth break within five years due to the bonding failure and need to be restored again [5]. The bonding durability is the most challenging issue in contemporary adhesive dentistry [6].

Extensive defects are found throughout the hybrid layer and interfacial failure accounts for 73.9% of the total restoration failures [7], which is attributed to the inadequate infiltration of adhesive into the DD matrix. Therefore, there is a common consensus that complete infiltration of the adhesive into the DD matrix is required for ideal dentin bonding [8]. In the last few decades, various approaches have been explored to promote adhesive’s infiltration in DD matrix and improve the bonding durability [9–12], for instance, upgrading dentin adhesive systems from 1st to 8th generation [13], replacing water with ethanol to increase adhesive infiltration [14–16], improving the polymerization degree of conversion and esterase resistance of the dentin adhesives [17, 18], inhibiting the activity of matrix metalloproteinases and cysteine cathepsins [19–21], improving infiltration under pressure assistance [22], remineralizing unwrapped collagens in the hybrid layer to improve their mechanical properties and resistance to collagenolytic enzymes [23], and etc. However, all these approaches improve the bonding durability slightly.

In nature, the hybrid layer is an interpenetrating network of biological composites comprised of 3-D DD matrix and polymerized adhesives. Inter-dispersion and interfacial compatibility of different substance phases are the two core scientific problems in the fields of composite materials engineering. The DD matrix is composed of >90 wt% collagen fibers and <10 wt% non-collagenous proteins (NCPs). Under physiological conditions, the NCPs rich in negative charges are highly-hydrated forming a hydrogel-like structure on collagen fibers [24]. Thus, there is a significant interfacial compatibility conflict between the hydrophobic adhesive molecules and the NCPs-covered collagen fibers. The infiltration of the hydrophobic molecules is severely limited by the hydrogel-like structure in the DD matrix. Moreover, once cured, micro-phase separation occurred extensively between the cured adhesive phase and the NCPs-covered collagen fibers. As a result, the hybrid layer is nonuniform and full of defects [25]. Therefore, we believe the inherent interfacial compatibility conflict between the hydrophobic adhesive molecules and the hydrophilic DD matrix is the main cause for the inadequate infiltration and bonding durability problem.

In this study, we hydrophobize the DD marix with octadecyltrichlorosilane (OTS) to reconcile the interfacial compatibility conflict between the DD matrix and the hydrophobic adhesive molecules, with the expectation of enabling the hydrophobic molecules to adequately infiltrate in the DD matrix and improving the bonding durability. Accordingly, three scientific tasks are undertaken in this study: (1) verify the interfacial compatibility conflict is the main cause for inadequate infiltration and bonding durability; (2) explore how reconciling interfacial compatibility conflict facilitates infiltration of the hydrophobic adhesive molecuels and improve bonding effectiveness and durability, and (3) confirm that resolving issues with interfacial compatibility conflict can overcome adhesive infiltration problem in the DD matrix and overcome bonding durability problem.

## 2. Experimental Section

### 2.1. Specimen preparation

Caries-free third molars extracted for orthodontic reasons or pericoronitis were collected with the individuals’ informed consent. The extracted teeth were placed in 0.1% NaN_3_ solution immediately after extraction, and used within one month after extraction. Crown enamel of each tooth was removed with a slow-speed Isomet saw (Buehler, Lake Bluff, IL, USA) under water-cooling. The surface of each dentin specimen was ground with 500-grift silicon carbide paper under running water in order to create a standardized smear layer [26]. Subsequently, specimens were randomly divided into three parts then treated as follows:

#### 2.1.1. Control group

Dentin in the Control group was prepared according to the typical wet-bonding technique. In brief, dentin surfaces were etched with 37% phosphoric acid (Aladdin, Shanghai, CHN) for 30 s, rinsed with deionized water for 1 min, and kept moist to prevent the collapse of the exposed organic dentin matrix (Samples were immersed in water until the next step, and taken out from water and blown for 1 to 2 seconds with an air syringe at a distance of about 2 cm to remove only the excess water).

#### 2.1.2. mEtOH group

The mEtOH group was prepared via a modified ethanol-wet bonding technique with selective removal of non-collagenous protein and a dehydration process stricter than the chemical dehydration protocol previously described by Sadek et al.[27]. In brief, after phosphoric acid etching, the samples were further treated with 5.25% sodium hypochlorite solution (NaClO, Aladdin, Shanghai, CHN) for 30 s to selectively remove the NCPs and sufficiently expose dentin collagen, then rinsed for 1 min with deionized water, and gradually dehydrated in ascending gradients of ethanol solution, containing 30%, 50%, 70%, 90%, and 100% ethanol, twice, for 10 min each.

#### 2.1.3. OTS group

In order for the demineralized dentin collagen to become hydrophobic, OTS was employed, one of the already clinically used silane coupling agents employed for the modification of ceramic or metal restorations [28]. In brief, after sequential treatment with phosphoric acid, sodium hypochlorite, and dehydration of gradient ethanol, samples were treated with 2% (v/v) OTS (Sigma-Aldrich, St. Louis, MO, USA) ethanol solution for 5 min at room temperature (Figure S1), and then immersed in 100% ethanol to remove un-reacted OTS.

### 2.2. Scanning electron microscopy

Prepared samples (*n*=3 each group) were sputter-coated with platinum and observed by field emission scanning electron microscopy (SEM, SU-70, Hitachi, JPN) at 3 kV through a secondary electron detector.

### 2.3. Contact angle measurement

The wettability of the dentin surfaces from three groups (*n* = 5 each group) was examined by the static contact angle measurement of 1 μL deionized water through a contact angle goniometer (CA 15plus, DataPhysics, Germany) using Visiodrop^®^ software. For each sample, three contact angle values were obtained at randomly selected sites without overlap. The values belonging to the same sample were averaged for statistical analysis. One-way ANOVA was used to compare the contact angles.

### 2.4. Fourier-transform infrared spectroscopy

Prepared samples were also evaluated for collagen integrity and OTS grafting by Fourier transform infrared spectroscopy (FTIR, Nicolet 6700, Thermo Fisher scientific LLC, USA). Dentin surface analysis was conducted over a spectral range of 4000-525 cm^-1^ with a smart Omni-sampler accessory at a resolution of 4 cm^-1^.

### 2.5. Protease digestion susceptibility

Dentin samples were cut into 5 mm × 3 mm × 1 mm pieces, and then prepared into the Control and OTS groups. Collagenase II and trypsin were used to test susceptibility to protease digestion.

Collagenase stock solution was prepared as follows: 100 mg collagenase II was dissolved in 3 mL tricine buffer solution (0.05 M Tricine buffer with 20 mM of CaCl_2_, pH 7.5) to prepare a stock-solution.The stock-solution was diluted 500 times with tricine buffer and used for the collagenase digestion experiment.

#### 2.5.1. Collagenase II digestion experiment

Each sample (*n*=5) was placed in a 2 mL centrifuge tube containing 320 μL of collagenase II (Gibco, Thermo Fisher Scientific Inc., MA, USA) solution at 37 °C. At 0, 0.5, 1, 1.5, 2, 3, 4, 6, 8, 12 and 24 h time points, 20 μL of solution was pipetted out and stored at −20°C. Hydrolyzed collagen was quantitatively determined by hydroxyproline in the solution with a hydroxyproline assay kit (BioVision Inc., CA, USA). Dentin samples (*n*=3) digested for 2 h and 8 h were also prepared for SEM observation. At each time-point, dentin samples were taken out of the collagenase solution, consecutively dehydrated in ascending gradients of ethanol, 30%, 50%, 70%, 90%, and 100% (each for 10 min), dried in hexamethyldisilazane (HMDS) twice (for 10 min each), sputter-coated with platinum and observed with SEM (SU-70, Hitachi, Japan).

#### 2.5.2. Trypsin digestion experiment

Trypsin stock solution contained 10 μg/mL trypsin (≥10000 BAEE, Sigma-Aldrich, St. Louis, MO, USA) in 0.01 M Tris-HCl buffer with 20 mM of CaCl2 (pH 7.0). Each test sample was placed in a 1.5 mL centrifuge tube containing 200 μL of trypsin stock solution and kept at 37 °C. At day 1 and day 7, samples (n=3 for each group, each time point) were drawn from the enzyme solution, dehydrated in ethanol, dried in HMDS as described before, and finally observed with SEM (SU-70, Hitachi, Japan).

### 2.6. Microtensile bonding strength evaluation

A hydrophobic adhesive, consisting of 59.5 wt% Bis-GMA, 39.5 wt% TEGDMA and 1 wt% camphorquinone(the three basic and critical hydrophobic components for all the dental adhesives, all purchased from Sigma) was produced in our lab according to an open formula of Heliobond (an enamel adhesive, IvoclarVivadent, Schaan, Liechtenstein) and referred to as “Hydrophobic-Adh” for the sake of simplicity. Prepared dentin surfaces in the Control, mEtOH and OTS groups were bonded with Prime&Bond NT (Dentisply, Konstanz, Germany), Optibond S (Keer, Orange, CA, USA) or Hydrophobic-Adh strictly following the instructions of the manufacturer (*n*=10 each group in each adhesive system). All samples were cured with a Radii Plus LED light-curing unit (SDI, Bayswater, Victoria, Australia, output of 1500 mWcm^-2^). To model restoration, resin composite layers were built up in 1-mm-thick increments to 3 mm thickness (Z250, 3M ESPE, St. Paul, MN, USA). After being stored in deionized water at 37°C for 24 h, samples were sectioned longitudinally into 1.0 mm ×1.0 mm × 8.0 mm beams by a slow-speed Isomet saw during water-cooling. 15 to 30 beams could be obtained from one tooth, depending on the size of the tooth. Beams were randomly divided into three equal sub-groups, one third of the samples were tested immediately, while the other two thirds were stored in artificial saliva [29] at 37 °C for 3 or 12 months in order to evaluate the durability of dentin bonding. Samples were stressed to failure under tension with a Micro Tensile Tester (Bisco, Schaumburg, IL, USA) at a crosshead speed of 1 mm/min. The force at failure was recorded, and the micro tensile bonding strength (MTBS) was calculated for each beam from the force at failure divided by the cross-sectional area of the fracture site. Each time, 5 to 10 beams from the same tooth were subjected to the micro-tensile bonding strength test, so 5 to 10 values could be obtained from the same tooth. Considering the random selection of samples prepared from the same tooth, a repeated analysis of variance was conducted to compare bonding strengths within each time point, and between time points within the same group.

### 2.7. Interfacial nanoleakage evaluation

Small water-rich and resin-poor regions within the polymerized hybrid layer (termed nanoleakage) can be identified using water-soluble tracers. In the present study, nanoleakage was evaluated by the uptake of ammonia complexed silver nitrate. The test solution was prepared as follows: 10 g AgNO_3_ was completely dissolved in the same mass of deionized water, then concentrated ammonia solution was added dropwise until the solution became translucent. The final concentration of AgNO_3_ in the ammonia-complexed silver solution was approximately 0.47 gmL^-1^.

From each tooth, two resin-bonded beams with a 1.0 mm ×1.0 mm cross-sectional area were randomly selected for interfacial nanoleakage evaluation. All beams were immersed into the AgNO_3_ test solution in the dark for 24 h, then rinsed thoroughly with distilled water, and immersed in photo developing solution for 8 h. Specimens were embedded in epoxy resin, polished with SiC paper and diamond paste, ultrasonically cleaned, and air-dried. The samples, after being coated with carbon, were analyzed by SEM (S-3700N, Hitachi, Japan) in the backscattered electron mode. For each sample, five randomly non-overlapping 1000× microscopic fields were photographed. From the micrographs, the total length of the dentin-adhesive interface and the length of interface with silver particles (representing nanoleakage) were measured with the UTHSCSA Image Tool 3.0 software. The extent of nanoleakage was calculated as the ratio of the length of interface with silver particles to the full length of each SEM photograph. Ten nanoleakage values obtained from two samples of the same tooth were averaged for statistical purpose. Independent-sample t-tests were used to compare nanoleakage values between groups at each time point, and pairwise t-tests were used to compare the nanoleakage between two time points within related sample groups.

### 2.8. Structure analysis

Structures of Bis-GMA, TEGDMA and Nile red were obtained from the PubChem database (CID 15284, http://pubchem.ncbi.nlm.nih.gov/compound/15284#section=Top and CID 7979, https://pubchem.ncbi.nlm.nih.gov/compound/7979#section=Top). VMD (visual molecular dynamics) was used to visualize the structure and measure the distance between atoms.

### 2.9. Infiltration study of hydrophobic monomers in demineralized dentin matrix

0.05 w/v% Nile red acetone solution was dissolved into the adhesive to dilute to 0. 01 w/v%, and the adhesives were applied on the prepared dentin surface from the OTS, mEtOH and Control groups. Samples were longitudinally cut into slices with a thickness of 1 mm and observed by confocal laser scanning microscopy (CLSM, Leica TCSSP8 MP, Leica microsystems, GER). Semi-quantitative analysis was conducted through the assay of the horizontal and vertical fluorescence intensity of the hybrid layer. The fluorescence density of the adhesive layer was set as 255, and the dentin layer was set as 0. The intensity of the hybrid layer was calculated from the horizontal fluorescence intensity with data over 200 eliminated, as regions with fluorescence intensity over 200 were considered as dentinal tubules.

### 2.10. Hybrid layer observation

Prepared dentin surfaces from the OTS, mEtOH and Control groups after Prime&Bond NT, Optibond S or Hydrophobic-Adh adhesives were applied were prepared. Small, rectangular sticks with end dimensions of approximately 0.6 mm by 1 mm were sectioned from the prepared sample in a direction perpendicular to the resin-dentin interface. 4 sticks from 2 teeth of each group were prepared. Further samples were prepared according to procedures used for TEM examination of calcified tissues [30]. Briefly, sticks were immersed in an aqueous ethylenediaminetetraacetic acid (EDTA) solution (41.3 gL^-1^ EDTA, pH 7.0) for 12 h at 4 °C. The partially decalcified specimens were rinsed in 2 mL of 0.1 M PBS (3×10 min) then fixed in 2.5% glutaraldehyde containing phosphate butter solution (PBS, 0.1 M, pH 7.0) at 4°C overnight. After washing with PBS for 3×10 min, specimens were post-fixated in 1% osmium tetroxide containing PBS (0.1 M, pH 7.4) for 3 h. Fixed samples were dehydrated in an ethanol gradient from 30% to 100%, and transferred into absolute acetone for 20 min. Samples were then infiltrated in a 1:1 mixture of Spurr resin and acetone for 1 h at RT, then treated with 3:1 resin/acetone for 3 h, and finally kept in absolute resin overnight. 90 nm thick ultrathin sections of were cut by means of a diamond knife (Diatome, Bienne, Switzerland) in an ultramicrotome (Sorvall Ultra Microtome MT5000, RMC, Tucson, AR) and positively stained with saturated uranyl acetate solution for 5 min followed by lead citrate solution (about 0.1 M) for 3 min. A JEOL TEM (1200 EX, Peabody, MA) was used to investigate the influence of OTS modification on dentin bonding.

## 3. Results and Discussion

### 3.1. Hydrophobic Modification of the DD collagen matrix to be compatible with hydrophobic adhesive monomers

In this study, a modified ethanol wet bonding technique featuring with a fully-exposed collagen matrix (Figure 1a) was developed by removing the highly-hydrophilic NCPs [31] and replacing inter-fibrous water using a strict ethanol gradient dehydration procedure. On this basis, collagen matrix was further modified with OTS, achieving a hydrophobic DD collagen matrix (Figure S1). Thus, a novel hydrophobic bonding strategy was developed to address the biggest challenging issue in adhesive dentistry. No detectable changes on the inherent structure of the dentin collagen fibers were found after the removal of NCPs and the modification of OTS (Figure 1a). The FTIR analysis showed the structural integrity of collagen and the successful modification of OTS as evidenced by the characteristic peaks at 1020 and 1080 cm^-1^ arising the OTS group, which can be attributed to C-O and Si-O vibrations, respectively (Figure 1b) [32]. The water contact angle of DD matrix increased from 66.8±1.9° in the Control group to 102.8±2.4° in the OTS group (p<0.05, Figure 1c). Moreover, the OTS modification greatly enhanced the resistance of the resulting collagen matrix to collagenase II (Figure 1d), one of the leading factors in the degradation of the collagen matrix in human dentin [33]. The degradation of collagen decreased by more than 90% compared with that of the Control group (Figure 1d3). The increased resistance of the OTS-modified collagen to proteases could be attributed to the change in surface properties of the collagen substrate, namely, the increased hydrophobicity and steric hindrance resulting from the grafting of long alkyl chains (-C_18_H_37_) on the collagen fibers. The protease-resistance of the OTS-modified collagen matrix may contribute to the stability of the resultant hybrid layer.

**Figure 1.**
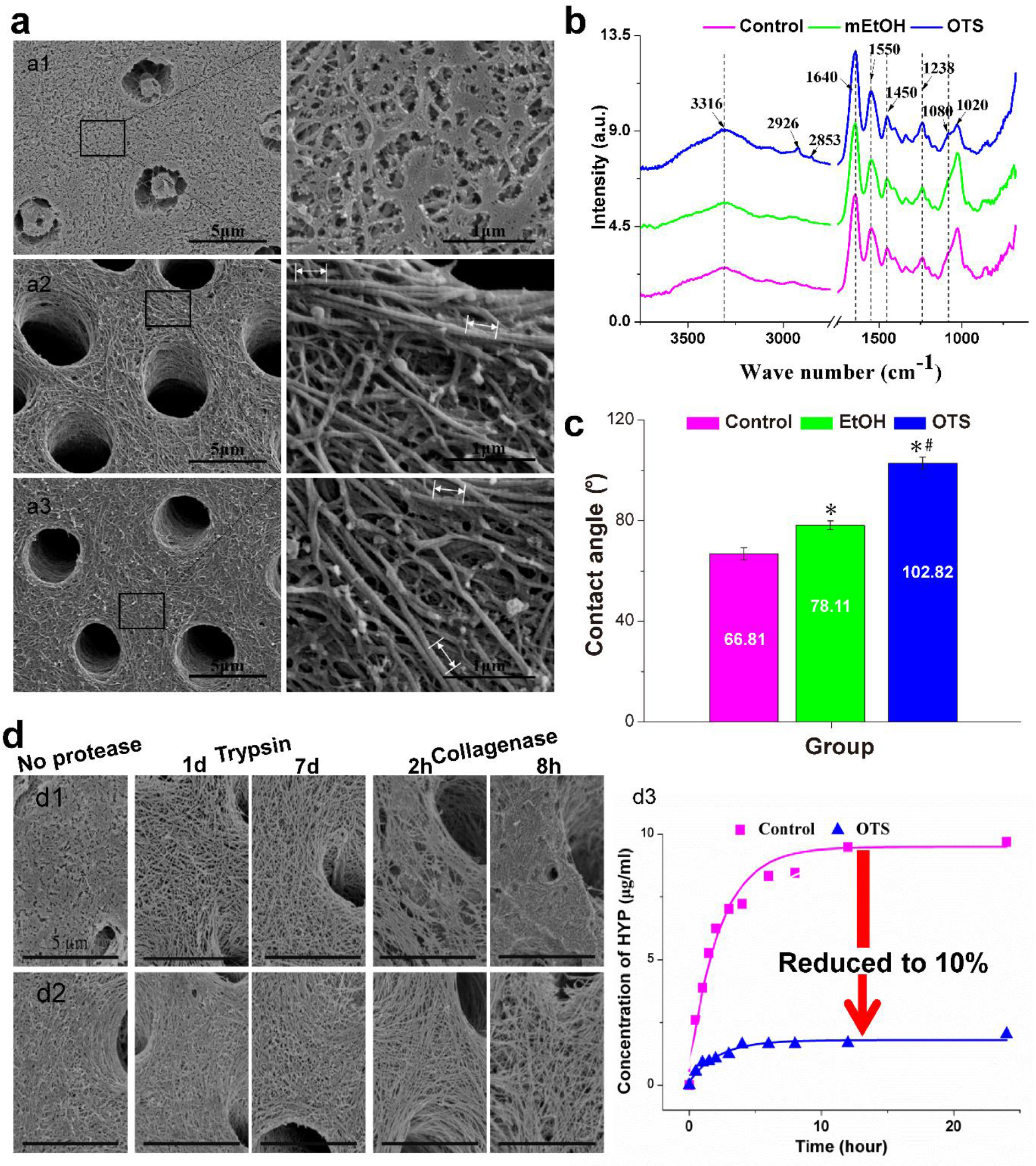
OTS modification changed the DD matrix from being hydrophilic to hydrophobic and greatly improved their resistance to proteinases without influencing the native structure of the collagen fibers. SEM images showed that after the demineralized dentin surface (a1, Control group) is treated with NaClO (a2, mEtOH group), the collagen network is sufficiently exposed with cross-bands of collagen fibers clearly visible (Note: length of five cross-bands is indicated by white arrows, with the length per cross-band calculated to be 67.2 nm). Notably, OTS modification has no detectable morphological influence on dentin collagen (a3, OTS group). FTIR spectra analysis (b) and contact angle measurements (c) indicate that OTS modification converts the hydrophilic dentin surface into a hydrophobic surface, with the collagen structure well preserved. SEM images of the dentin surfaces before and after being exposed to trypsin or collagenase II (the Control group, d1, and the OTS group, d2) and quantitative determination of dentin collagen hydrolyzed by collagenase II (d3), show that the OTS-modified collagen matrix possesses improved protease-resistance.

### 3.2. Reconciling interfacial compatibility conflict greatly increases bonding effectiveness

NCPs are believed to be primarily responsible for the controlled growth of the mineral phase involved in biomineralization [34]. However, NCPs are considered to be the greatest interference factor in adhesive infiltration in dentin bonding as well as interfibrous water. For the Control group, the during 12-month aging, the bonding strength decreased from 33.80 MPa to 18.0 MPa in the Optibond S system (with an attenuation rate of 5.11% per month) and from 22.53 MPa to 6.7 MPa in the Prime&Bond NT (with an attenuation rate of 9.99% per month). By eliminating the two interferences, according to the existing knowledge of adhesive theory, theoretically, complete infiltration of dentin adhesives into the DD matrix with a flawless hybrid layer and an ideal bonding effectiveness and durability can be achieved in the mEtOH groups. Actually, the immediate bonding strength increased in both bonding systems (33.80 to 35.09 in the Optibond S system, 22.53 MPa to 30.63 MPa in the Prime&Bond NT system, Figure 2a). However, silver particles still accumulate throughout the whole hybrid layer even immediately after bonding, indicating extensive nanoleakage in the hybrid layer (Figure 2b, c). Moreover, during 12-month aging, the bonding strength decreased rapidly as the nanoleakage increased, and the bonding strength decreased 31.80% in the Optibond S system (35.09 MPa to 23.93 Mpa, with an attenuation rate of 3.14% per month) and 32.81 % in the Prime&Bond NT (30.63 MPa to 20.58 MPa, with an attenuation rate of 3.26% per month). These results suggest that without the interferences of NPCs and water, extensive defects still produce throughout the hybrid layer immediately after bonding, and the bonding strength decreases rapidly over a 12-month period.

**Figure 2.**
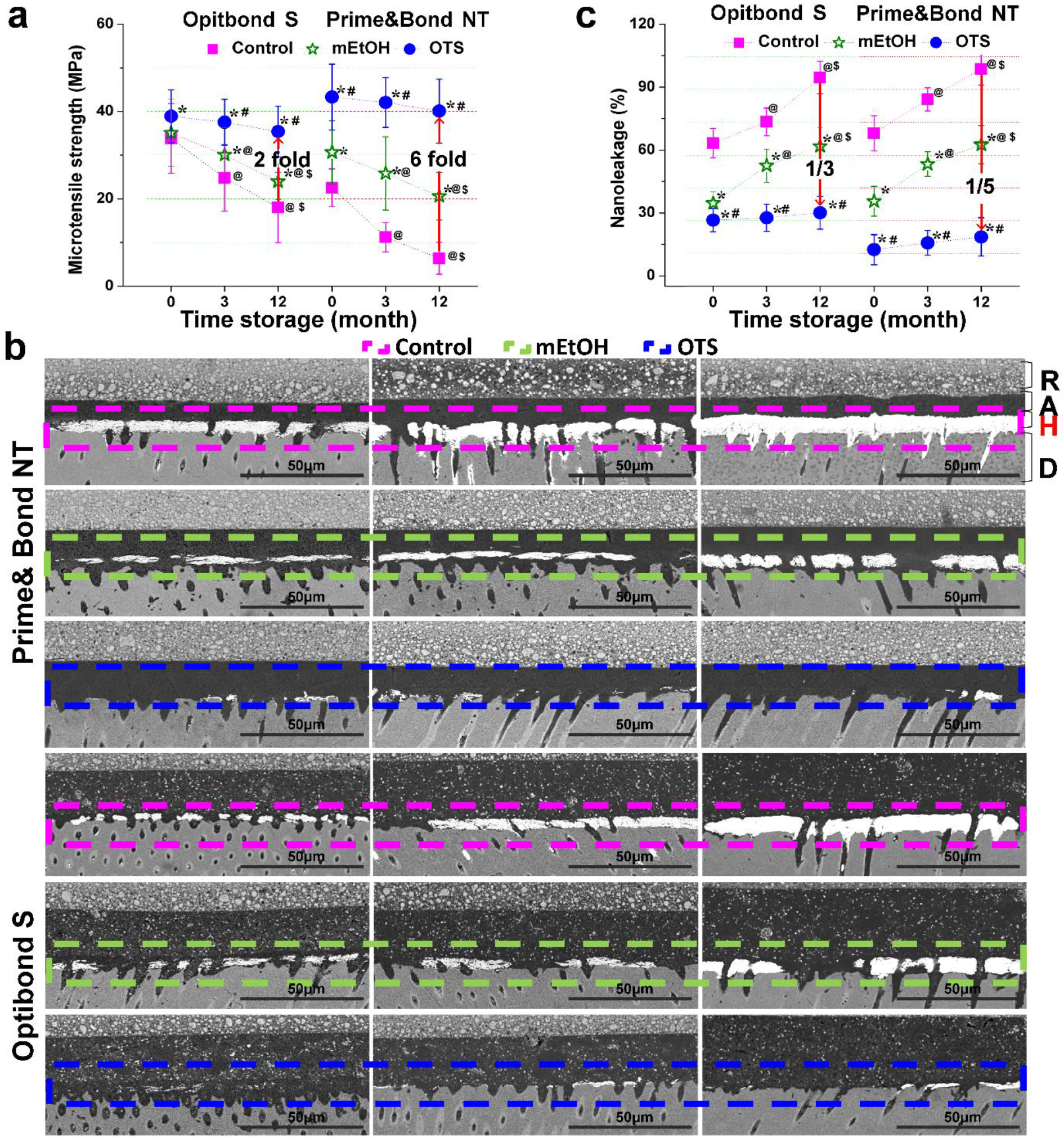
Reconciling the interfacial compatibility conflict between the hydrophobized DD matrix and the hydrophobic adhesive molecules greatly improve bonding effectiveness and durability. Microtensile bonding strength analysis (a) and nanoleakage analysis (b and c) show that modulation of the interfacial compatibility between the demineralized dentin collagen and the critical adhesive monomers (OTS group) significantly improves dentin bonding effectiveness and durability. Hybrid layers produced by the novel bonding strategy show much less nanoleakage than those produced by wet bonding for both of the dentin adhesives applied. * p < 0.05 vs. the Control group at the same time point. # p < 0.05 vs. the mEtOH group at the same time point. @ p < 0.05 vs. the immediate test index of the same group. R: resin; A: adhesive layer; H: hybrid layer; D: dentin. Scale bar: 50 μm.

Conversely, the OTS groups achieved much higher immediate bonding strength in either the Optibond S or Prime&Bond NT system, being 1.15 times in the Opitibond S system and 1.92 times in the Prime&Bond NT system compared with the Control groups (33.8 MPa to 38.9 MPa in the Optibond S system, 22.5 MPa up to 43.2 MPa in the Prime&Bond NT system). Moreover, the attenuation rate of the bonding strength during 12-month aging was much smaller (0.64% per month decrease in the Optibond S system and 0.78% per month decrease in the Prime&Bond NT, Table 1), achieving a bonding strength1.7-2 times that of the mEtOH groups (35.4 MPa vs 23.93 MPa in the Optibond S system, and 40 Mpa vs 20.58 MPa in the Prime&Bond NT system) and 2-6 times that of the Control groups (35.4 MPa vs 18.0 MPa in the Optibond S system, and 40 Mpa vs 6.7 MPa in the Prime&Bond NT system). The hybrid layers of the OTS groups also demonstrated a greatly reduced nanoleakage (Figure 2b, c). These results indicate that reconciling the interfacial compatibility conflict between the DD matrix and the critical hydrophobic adhesive monomers leads to the formation of a more homogeneous hybrid layer with a markedly reduced number of defects, which in turn greatly improves bonding strength and durability.

**Table 1.** Analysis of attenuationin microtensile bonding strength

| Adhesive | Group | Microtensile strength reduced ( % ) |  |  |  |
| --- | --- | --- | --- | --- | --- |
|  |  | First 3 months |  | 12 months |  |
|  |  | total | per month | total | per month |
| Prime & Bond NT | Control | 50.30 | 20.79 | 71.73 | 9.99 |
|  | mEtOH | 15.77 | 5.56 | 32.81 | 3.26 |
|  | OTS | 2.87 | 0.96 | 7.43 | 0.64 |
| Optibond S | Control | 26.66 | 9.82 | 46.75 | 5.11 |
|  | mEtOH | 14.35 | 5.03 | 31.80 | 3.14 |
|  | OTS | 3.48 | 1.17 | 9.01 | 0.78 |
| Hydrophobic-Adh | mEtOH | 16.31 | 5.76 | 33.25 | 3.31 |
|  | OTS | 0.01 | 0.00 | 0.12 | 0.01 |

### 3.3. High interfacial compatibility produces ideal bonding effectiveness

To assist the infiltration of the critical hydrophobic adhesive monomers into the highly hydrated DD matrix, solvent and a high number of amphiphilic monomers are incorporated in commercial dentin adhesives. However, in this present study, for the mEtOH groups and the OTS groups, the hydrophobic adhesive components can infiltrate the DD matrix without the help of the solvent or amphiphilic monomers. Instead, the presence of them negatively influences the rate and efficiency of polymerization, produces hydrophilic micro-regions, and simultaneously increases water affinity at the resin-dentin interface, which is detrimental to bonding strength and durability[35]. To further investigate the direct interaction between the critical hydrophobic monomers and hydrophobized demineralized collagen matrix, one hydrophobic adhesive, Hydrophobic-Adh, consisting only of the critical hydrophobic adhesive components Bis-GMA, TEGDMA and CQ, was used.

The bonding effectiveness and durability of the mEtOH group bonded with Hydrophobic-Adv was improved compared with the two corresponding mEtOH groups bonded with Prime&Bond NT and Optibond S (Figure 3a). However, extensive defects were still found throughout the hybrid layer immediately after bonding (Figure 3b). During the 12-month aging, the bonding strength decreased to 66.74% of the immediate bonding strength (from 38.53 MPa to 25.72 MPa, with an attenuation rate of 3.31% per month) with nanoleakage increasing rapidly. These results confirm that the interfacial compatibility conflict between the critical hydrophobic monomers and the DD collagen matrix still result in extensive defects throughout the hybrid layer. In contrast, for the OTS group using Hydrophobic-Adv, observation of silver accumulation showed a flawless hybrid layer (Figure 3b). The aging study reveals that no attenuation occurred in bonding strength during the whole observation period (from 46.83 MPa to 46.77MPa, with an attenuation rate of 0.01% per month), showing incredible stability. The 12-month bonding strength increased 7.3 fold compared with the Control group using Prime&Bond NT (Figure 2a), which was even higher than the immediate bonding strengths of the OTS groups using the two dentin adhesives. The 12-month nanoleakage was far lower than the immediate nanoleakage of the OTS group with the Prime&Bond NT system. Clearly, high interfacial compatibility between the hydrophobic DD matrix and the critical hydrophobic adhesive molecules produces a flawless hybrid layer and provides an ideal bonding effectiveness and durability.

**Figure 3.**
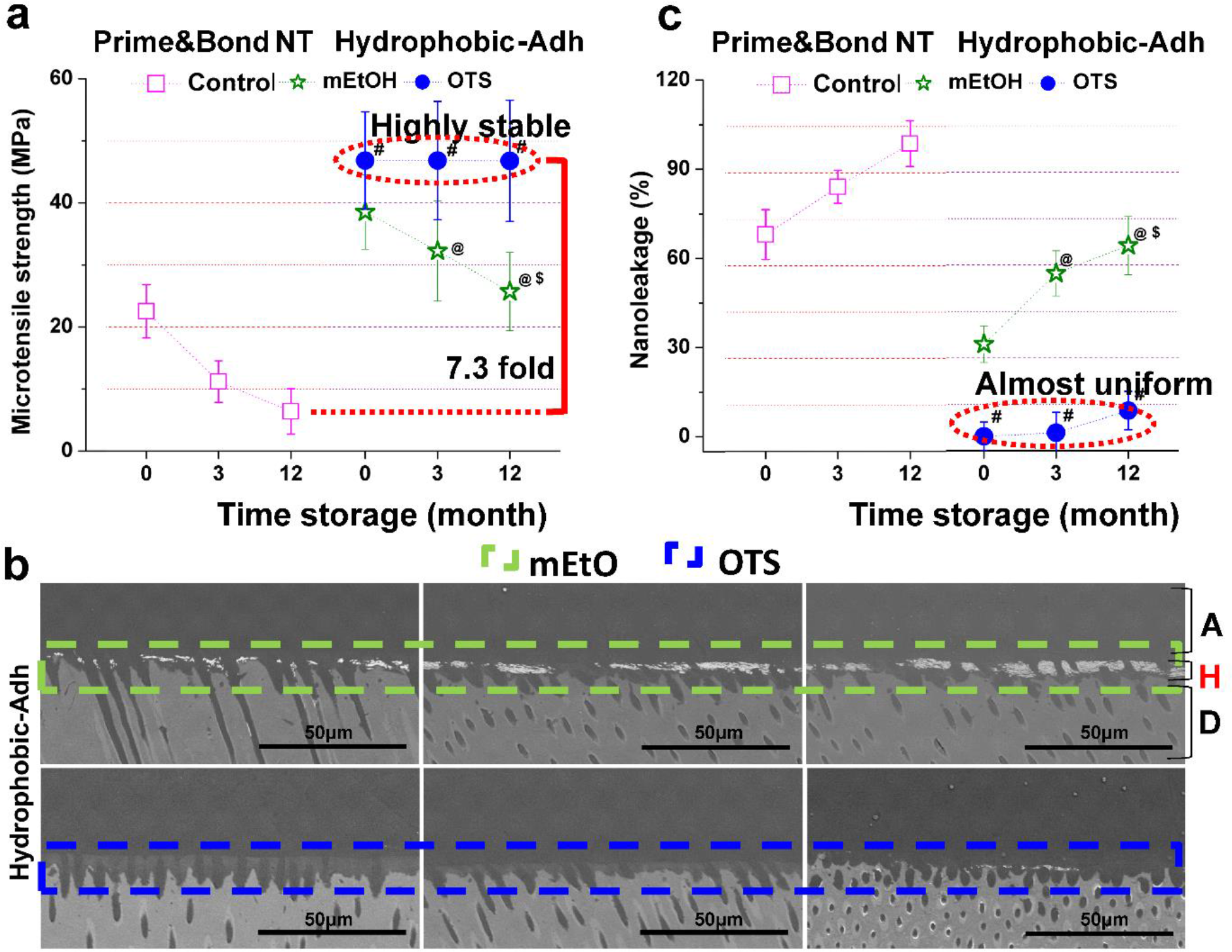
The Hydrophobic-Adv and the hydrophobic DD matrix formed a flawless hybrid layer and achieved ideal bonding effectiveness and durability. (a) Microtensile bonding strength evaluation. (b) Backscattered electron images and (c) semiquantitative analysis of nanoleakage analysis. # p < 0.05 vs. the mEtOH group at the same time point. @ p < 0.05 vs. the immediate test index of the same group. R: resin; A: adhesive layer; H: hybrid layer; D: dentin. Note: As the Control group is unsuitable for clinical dentin bonding with enamel adhesive based on wet bonding, it was not included in the Hydrophobic-Adh system (Figure S3).

### 3.4. Reconciling interfacial compatibility conflict facilitates adequate infiltration of the hydrophobic adhesive molecules into the DD matrix

To investigate how reconciling interfacial compatibility conflict between the DD matrix and the hydrophobic adhesive molecules influences the infiltration of the hydrophobic adhesive molecules in the DD matrix, tracing the relative content and distribution of the critical adhesive hydrophobic molecules in the DD matrix is of great importance. Herein, Nile red, a fluorescent molecule usually applied as a model hydrophobic molecule in in vitro cell-uptake testing and intracellular lipid droplet staining, is of comparable size and hydrophobicity with the typical hydrophobic adhesive monomers, Bis-GMA and TEGDMA (the distances between certain carbon atoms for Bis-GMA, Nile red and TEGDMA are 12.87 Å, 10.59 Å and 9.89 Å, respectively, Figure 4a). Therefore, the Nile red fluorescence intensity and distribution are positively correlated with the relative content and distribution of the critical hydrophobic adhesive monomers in the DD matrix.

**Figure 4.**
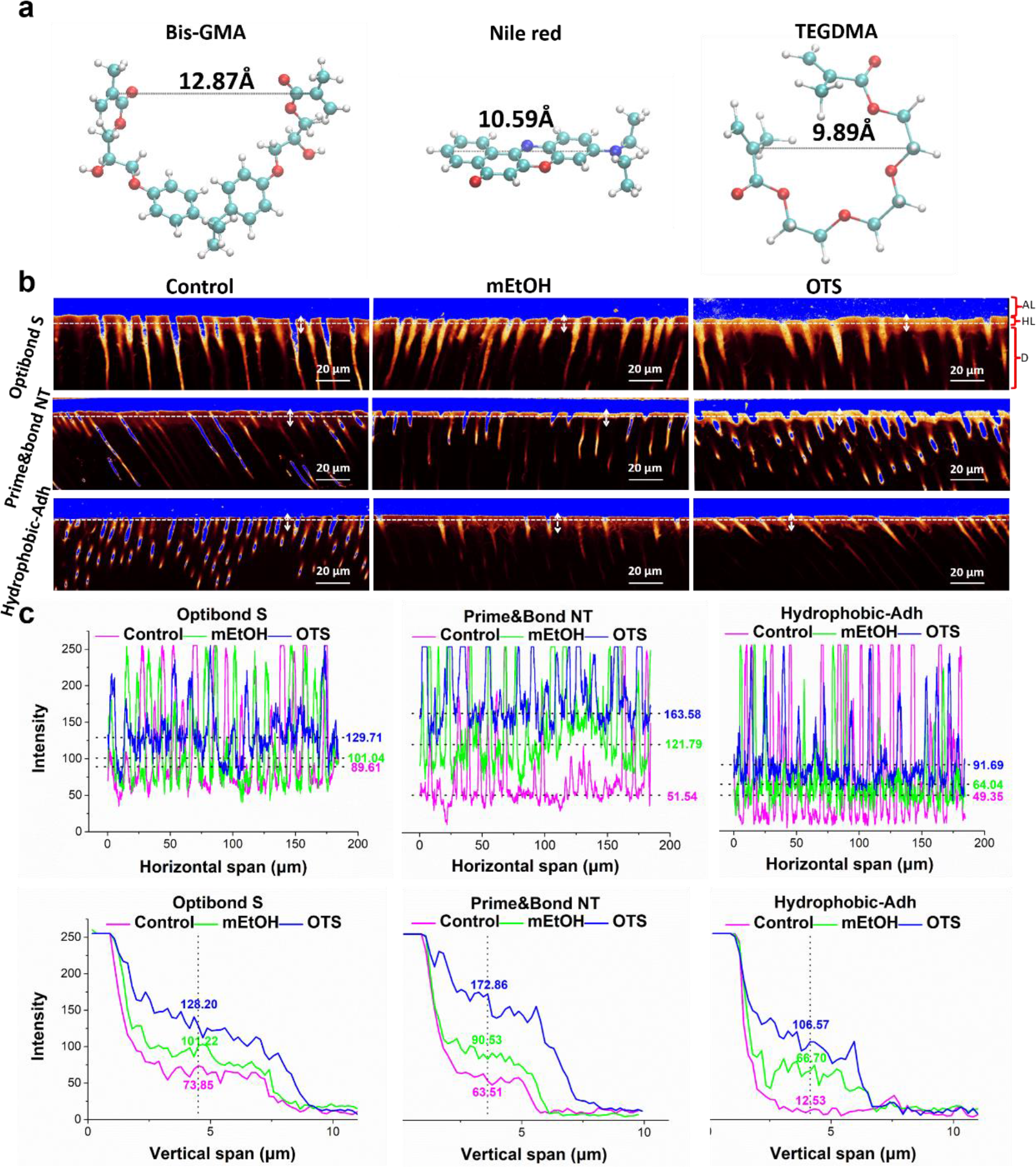
The interfacial compatibility determines the infiltration, relative content and distribution of the critical hydrophobic adhesive components in DD matrix. (a)Structure diagrams of two critical hydrophobic adhesive monomers and Nile red. (b)Images of confocal laser scanning microscopy for hydrophobic monomer infiltration traced with Nile red. (c) Semi-quantitative analysis of the fluorescence intensity (horizontal dotted line, indicate the middle layer of the hybrid layer; the vertical two-way arrow indicates the longitudinal transition from the adhesive layer to the hybrid layer and the dentin).

Due to the different distributions of hydrophobic Nile red, the adhesive layer is blue, the hybrid layer is red and the dentin layer is black in the overexposed confocal laser scanning microscopy image (Figure 4b). The fluorescence intensity of the hybrid layers in the OTS groups is the strongest, followed by the mEtOH groups and the Control groups, regardless of which type of adhesive was used. Moreover, the hybrid layers in the OTS groups also demonstrates maximum uniformity of fluorescence intensity. The results reveal an interesting but ignored fact that, although the wet bonding technique allows for infiltration of dentin adhesives into the DD matrix, the severe interfacial compatibility conflict between the DD matrix and the critical hydrophobic adhesive molecules lead to inadequate infiltration, nonuniform distribution and low relative content of the critical hydrophobic adhesive molecules in the hybrid layer. The Nile red tracer study clearly shows that reconciling the interfacial compatibility conflict of the two markedly promotes infiltration, distribution and the relative content of the hydrophobic components in the DD matrix. The highest interfacial compatibility between the hydrophobic DD matrix and the hydrophobic adhesive exhibited the most adequate infiltration, the highest relative content and the most uniform distribution.

### 3.5. Schematic bonding mechanism conditioned by the interfacial compatibility

Based on the abovementioned results, the bonding mechanism is graphically illustrated in Scheme 1. Once the dentin adhesive is applied onto the DD matrix, the adhesive is automatically divided into two layers: one part rapidly infiltrates into the DD matrix to form hybrid layer, the other remains above the DD matrix to form an adhesive layer (Scheme 1a1). For the Control groups, the water-full 3-D DD matrix acts as a strongly polar “NCPs modified collagen chromatography column”. According to the “like dissolves like” principle, the infiltration of the polar solvent and the amphiphilic monomers into the hydrated DD matrix is preferential. However, the entry of critical hydrophobic molecules is selectively limited. This inevitably causes the severe redistribution of the critical hydrophobic molecules between the DD matrix layer and the adhesive layer. Thus, more hydrophobic monomers exist in the adhesive layer (Scheme 1b11). This is also the reason for the lowest relative content of the hydrophobic components in the hybrid layer for the Control groups. Moreover, even in the DD matrix layer, the hydrophilic collagen fibers covered with hydrated NCPs preferentially absorb amphiphilic monomers and repel the critical hydrophobic molecules, producing a descending gradient distribution of the critical hydrophobic monomers around the collagen fibers (Scheme 1b12). Once polymerized, severe and extensive phase separation between the demineralized dentin collagen matrix and polymerized adhesive phase inevitably occurs, which produces numerous defects throughout the hybrid layer (Scheme 1, Figure 2b).

**Scheme 1.**
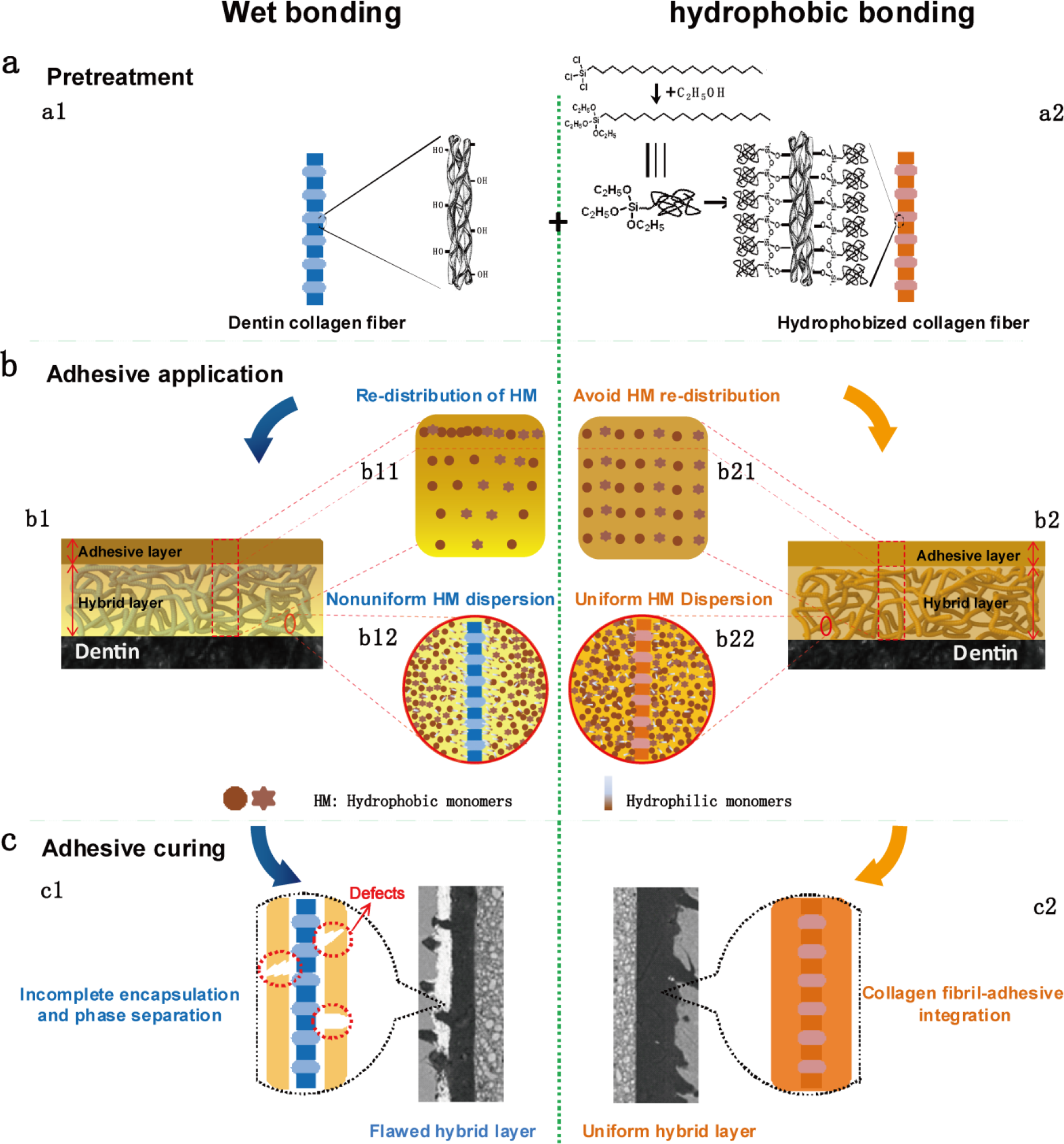
The scheme illustrates how the interfacial compatibility influences the infiltration and distribution of the hydrophobic molecules in the DD matrix and the final bonding effectiveness. The hydrophilic dentin collagen matrix in wet-bonding is represented by the blue schematic collagen fiber-like network (a1). The hydrophobized collagen matrix modified by OTS is represented by a yellow-colored network (a2). Once the adhesive is applied, redistribution of the critical hydrophobic monomers between the DD matrix and adhesive layers occurs, and the hydrophobic components preferentially distribute in the adhesive layer (b11). Hydrophilic collagen fibers repel hydrophobic monomers and preferentially adsorb amphiphilic monomers (b12), which results in extensive phase separation between the NCPs covered collagen fibers and cured adhesive, and inevitably produces numerous defects throughout the hybrid layer (c1). In contrast, for the hydrophobic dentin matrix, redistribution of the critical hydrophobic monomers between the DD matrix layer and the adhesive layer is avoided (b21). The hydrophobic collagen fibers can indiscriminately absorb the hydrophobic monomers and the amphiphilic monomers simultaneously (b22) to form a highly-integrated collagen-adhesive hybrid layer (c2).

For the mEtOH groups, because the two biggest interference factors, NCPs and interfibrous water, were removed, it was theoretically easy to obtain complete infiltration. However, due to the hydrophilicity of collagen fibers, the 3-D dentin collagen matrix layer still acts as a polar “Collagen chromatography column” and finally leads to the redistribution of the critical hydrophobic molecules between the DD matrix layer and the adhesive layer. The hydrophobic components still preferentially distribute in the adhesive layer, which is also verified by analysis of the fluorescence intensity (Figure 4b). In the DD matrix layer, the hydrophilic collagen fibers also preferentially absorb amphiphilic monomers and repel the critical hydrophobic adhesive monomers, producing a descending gradient distribution of the critical hydrophobic monomers around the collagen fibers. Following polymerization, phase separation between the polymerized adhesive and hydrophilic collagen fibers still occurs, producing extensive defects throughout the hybrid layer (Figure 2b and 3b).

In contrast, for the OTS groups, the hydrophobized DD matrix is compatible with the critical hydrophobic adhesive molecules (Scheme 1a2). The good interfacial compatibility avoids redistribution of the hydrophobic adhesive molecules between the DD collagen matrix layer and the adhesive layer and promote the infiltration, distribution and relative content of the hydrophobic molecules in the DD matrix (Scheme 1b21). This is confirmed by the strongest fluorescence intensity of the hybrid layer being observed in the OTS groups (Figure 4b). Moreover, the hydrophobized demineralized collagen fibers indiscriminately absorb the hydrophobic adhesive molecules or the amphiphilic molecules simultaneously (Scheme 1b22). Once polymerized, a hybrid layer is formed with markedly reduced defects (Scheme 1c2, Figure 2b). In particular, when the hydrophobic adhesive is applied on the hydrophobic DD matrix, the high interfacial compatibility allows for adequate infiltration and uniform distribution of adhesive molecules in the DD matrix, forming a flawless hybrid layer and producing ideal bonding effectiveness and durability (Figure 3).

### 3.6. Interfacial characteristics of a highly-integrated hybrid layer

To intuitively observe the microstructure of the hybrid layer particularly the collagen fiber-adhesive interface phase, TEM was used. Collagen fiber is composed of 300 nm long staggered collagen molecules spaced in multiples of D-periods (~67 nm) [36], a 40 nm gap region (0.6 D) plus a 27 nm overlap region (0.4 D, Figure 5). For the OTS group, due to the good interfacial compatibility, the critical hydrophobic adhesive monomers, with a size of approximately 1 nm, were uniformly distributed and randomly adsorbed in both the overlap region (~27 nm) and gap region (~40 nm), and even penetrated the inner layer of the hydrophobized collagen fibers (Figure 5d). *Once polymerized, a highly integrated collagen fiber-adhesive interface phase is formed, similar to the integration between the DD matrix and inorganic phase in mineralized dentin.* The highly integrated collagen fiber-adhesive interface phase weakens the difference in the visual boundary between the collagen fibers and the polymerized adhesive phase, and prevents polar reactive sites from reacting with uranyl acetate and lead citrate. This results in an indistinct collagen fiber-adhesive interface and even obscures the cross-bands of collagen fibers in the three OTS groups (Figure 5a-c, S3).

**Figure 5.**
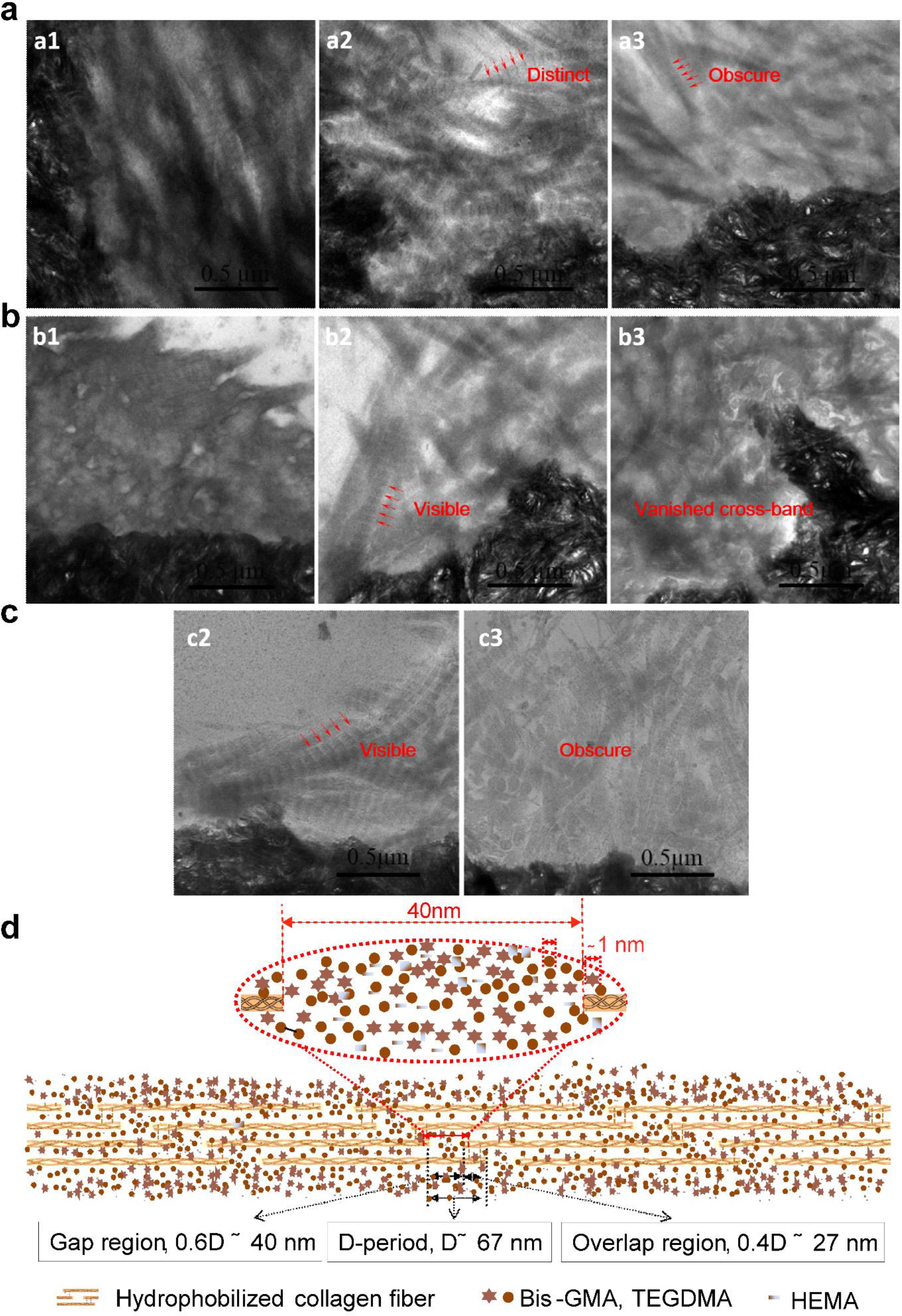
High interfacial compatibility produced a highly-integrated collagen fiber-adhesive hybrid layer. Typical TEM photomicrographs of hybrid layers in Optibond S (a), Prime&Bond NT (b) and Hydrophobic-Adh systems (c) from the Control (a1, b1), mEtOH (a2, b2 and c2) and OTS (a3, b3, and c3) groups. The critical hydrophobic adhesive monomers are indiscriminately absorbed onto both the overlap region (~27 nm) and gap region (~40 nm) of the OTS modified collagen fibers, and even penetrate the inner collagen fibers (d). Note: The dentin-adhesive interface of the Control group bonded with Hydrophobic-Adh was too weak to withstand the sectioning of the rectangular sticks for TEM observation, so no samples were obtained from this group, thus a C1 image is not present.

## 4. Conclusion

In conclusion, this study demonstrates that: (1) the interfacial compatibility conflict between the hydrophilic DD matrix and the hydrophobic adhesive molecules is the fundamental reason for the inadequate adhesive infiltration and the bonding durability problem; (2) reconciling the interfacial compatibility conflict markedly promote infiltration, relative content and uniform distribution of the critical hydrophobic adhesive molecules in the DD matrix, greatly enhancing bonding effectiveness and durability; (3) highly interfacial compatibility produces an almost flawless hybrid layer and ideal bonding strength and durability. This study clarifies the main cause of poor wet-bonding durability and demonstrates a paradigm in adhesive dentistry to overcome the long-existing bonding durability problem associated with inadequate adhesive infiltration into the DD matrix. This provides a new angle of view to resolve clinical dentin bonding durability problem and will significantly promote the adhesive dentistry.

## Supporting Information

Supporting Information is available online or from the author.

## Acknowledgements

The authors wish to thank Senior engineer Jingping Zhu for her help in SEM study.Thank Prof. Y. Q. Shen for his technical support and revising the manuscript. Thank Dr. Zexin Chen for his help in data analysis. Thank Dr. Yu Kang for her help in molecular structure analysis. The authors also wish to thank Prof. Hanying Li for their suggestions for manuscript revision. We all sincerely thank deceased Prof. Hui Chen for her help in the study.

## Fund

This study was supported by the National Natural Key R&D Program of China (2018YFC1105302) and the National Natural Scientific Funding (31670971, 81600835). Samples were acquired with the approval from the Ethics Committee of the Affiliated Stomatologic Hospital of the College of Medicine, Zhejiang University.

## Author contributions

Q. Luo and Y. Chen carried out the experiments and participated in analyzing the experimental data and writing the manuscript; J. Xu, C. Shu and Z. Li prepared the adhesive of “Hydrophobic-Adh” and carried out some of the experiments based on this adhesive; Y. Shen provided advice on paper writing, composition of the TOC and revised the manuscript. X. Li conceived the project, designed the experiments, analyzed the experimental data, and wrote the manuscript with contributions from all authors.

