## supplemental materials for "Reconciling Inherent Interfacial Compatibility Conflict Enhances Adhesive Infiltration and Resolves Dentin Bonding Durability"

FigureS1. Solvent selection.

Figure S2. Failure model analysis.

FigureS3. TEM observation on the influence of OTS modification on the appearance of collagen fibers.

1. **Solvent selection.** Before selection of ethanol as the solvent for OTS, experiments were carried out to evaluate three candidate solvents, namely ethanol, diethyl ether and toluene. A group modified with 2% (v/v) OTS ethanol solution for 5 min was named the OTS-E group, that with 0.1% (v/v) OTS diethyl ether solution for 60 s was named the OTS-D group, and that with 0.01% (v/v) OTS toluene solution for 5 s was named the OTS-T group. Surface morphology was observed by SEM after ethanol dehydration. Surface wettability was characterized by the contact angle. Bonding effectiveness and durability were assessed through immediate bonding strength after bonding with Prime&Bond NT.

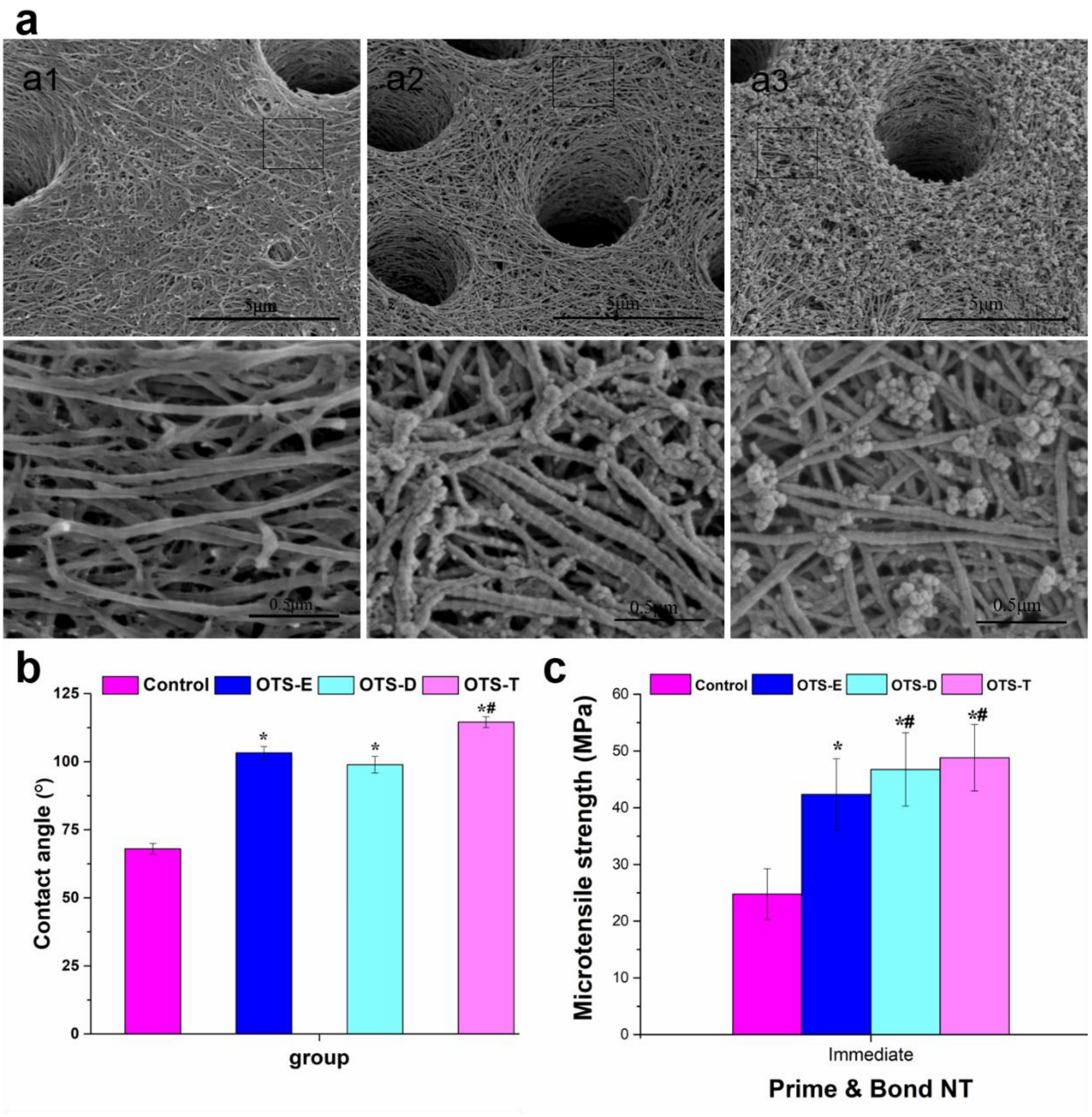

**FigureS1. Solvent selection.** Typical SEM images for OTS-E (a1), OTS-D (a2) and OTS-T groups (a3). All OTS-modified groups show an increase in contact angle values (b) and enhanced bonding strength (c). As the reaction speed was too fast to control for toluene, the ignition point was too low too safe use for diethyl ether, and the influence on the collagen morphology of the above two was too strong, ethanol was selected as the solvent for further systematic studies, which has the benefits of being non-toxic and safe to handle. * $p < 0.05$ vs. the Control group at the same time; $^{\#}$ $p < 0.05$ vs. the OTS-E group at the same time; $^{@}$ $p < 0.05$ vs. immediate microtensile strength of the same group.

2. **Typical failure mode.** The cross-sectional area of the fracture site after the MTBS test was verified by a digital caliper and classified as cohesive (failure exclusively within dentin or resin), adhesive (failure at the resin/dentin interface), mixed (failure at the resin/dentin interface that includes cohesive failure of the neighboring substrates), or premature failure (failure before the MTBS test). Specimens with premature failures were assigned to the tooth as 0 MPa, while those with cohesive failures were excluded. The constituent ratio of the failure model in each group was analyzed with the chi-square test. Typical failure models were observed with SEM (S3700, Hitachi, Japan) after sputter-coating with platinum.

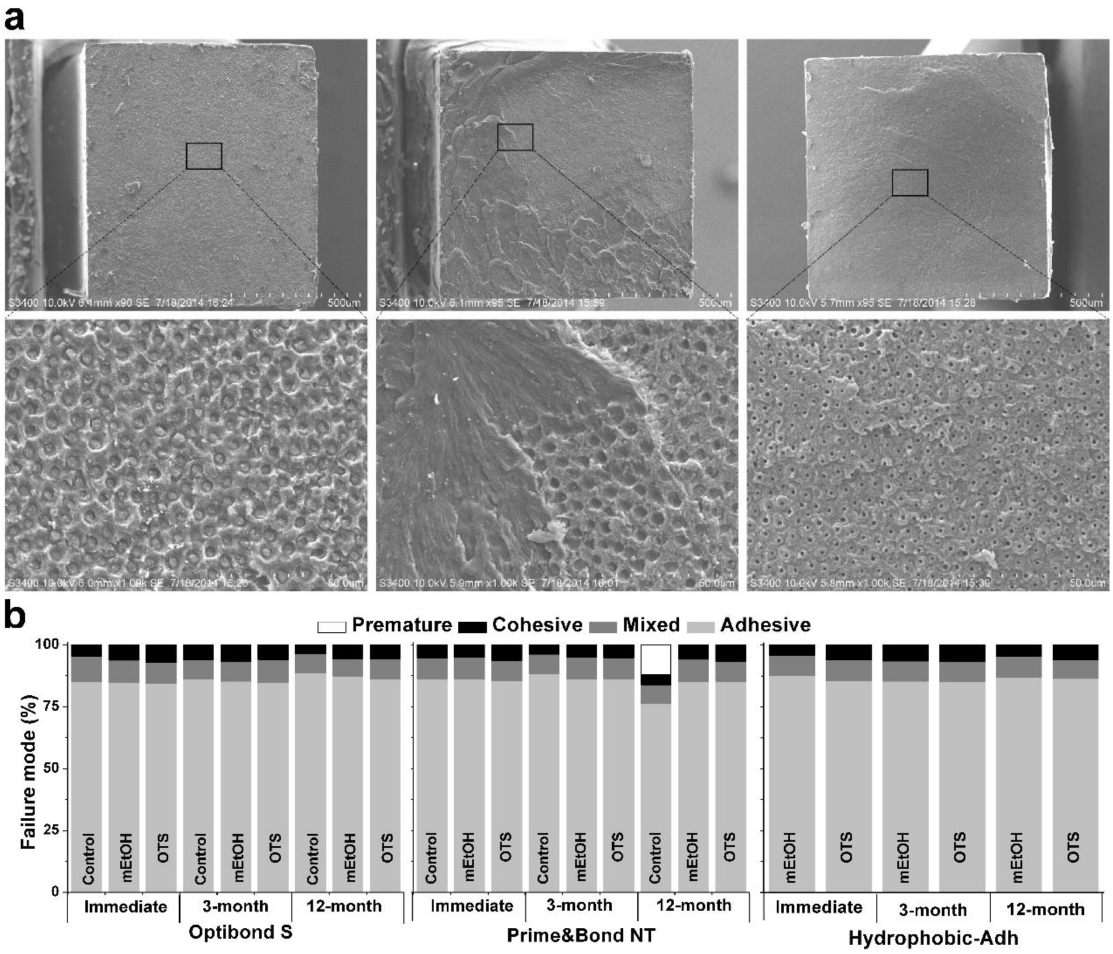


**Figure S2. Failure model analysis.** (a) SEM images of typical failure modes: Adhesive failure (failure at the resin/dentin interface, left column), mixed failure (failure at the resin/dentin interface that includes cohesive failure of the neighboring substrates, middle column) and cohesive failure (failure exclusively within dentin or resin, right column). (b) Failure mode analysis of Optibond S, Prime&BondNTand Hydrophobic-Adh systems. Most failure occurs at the resin/dentin interface. Premature failure is defined as failure before the MTBS test, and only appeared in the 12-month Control group.

3. **TEM observation on the influence of OTS modification on the appearance of collagen fibers.** Dentin surfaces from the Control group and the OTS group without the application of dentin adhesives were prepared for TEM observation.

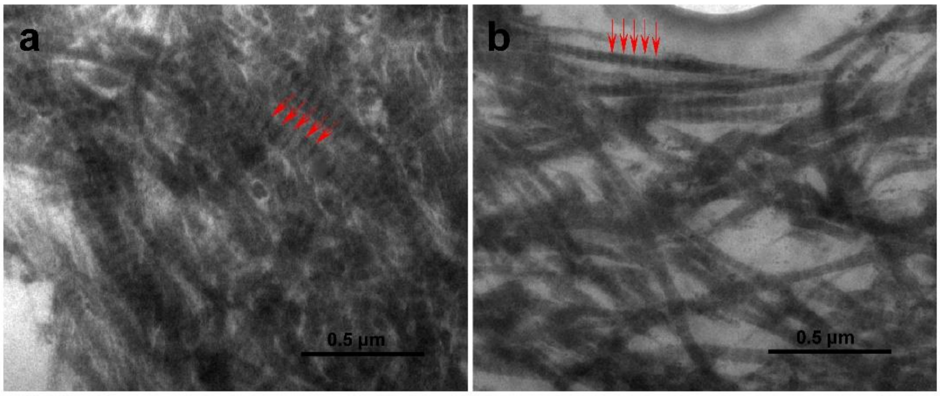


**Figure S3. OTS modification showscollagenfibers with similar appearanceunder TEM observation.** Typical TEM photomicrographs of collagen in the Control (a) and OTS groups (b) before the adhesive is applied. Collagen fibers of the Control group and the OTS group both show typical periodic cross-bands of collagen fibers.Collagen fibers are rich in polar amino acids, especially in the gap region,[17] which can induce the nucleation of uranyl acetate and lead citrate to produce cross bands visible in TEM. When OTS was chemically introduced into the collagen, the number of hydroxyl groups was greatly reduced, particularly in the gap region. However, a some charged groups on the polar amino acid residues were still present, and cross-bands of the collagen fibers were still stained clearly and distinctly, similar to that of the Control groups.
